# Exploiting collateral sensitivity controls growth of mixed culture of sensitive and resistant cells and decreases selection for resistant cells

**DOI:** 10.1101/2020.05.07.082073

**Authors:** Vince Kornél Grolmusz, Jinfeng Chen, Rena Emond, Patrick A. Cosgrove, Lance Pflieger, Aritro Nath, Philip J. Moos, Andrea H. Bild

## Abstract

**Background:** CDK4/6 inhibitors such as ribociclib are becoming widely used targeted therapies in hormone-receptor-positive (HR+) human epidermal growth factor receptor 2-negative (HER2-) breast cancer. However, cancers can advance due to drug resistance, a problem in which tumor heterogeneity and evolution are key features.

**Methods:** Ribociclib-resistant HR+/HER2-CAMA-1 breast cancer cells were generated through long-term ribociclib treatment. Characterization of sensitive and resistant cells were performed using RNA sequencing and whole exome sequencing. Lentiviral labeling with different fluorescent proteins enabled us to track the proliferation of sensitive and resistant cells under different treatments in a heterogeneous, 3D spheroid coculture system using imaging microscopy and flow cytometry.

**Results:** Transcriptional profiling of sensitive and resistant cells revealed the downregulation of the G2/M checkpoint in the resistant cells. Exploiting this acquired vulnerability; resistant cells exhibited collateral sensitivity for the Wee-1 inhibitor, adavosertib (AZD1775). The combination of ribociclib and adavosertib achieved additional antiproliferative effect exclusively in the cocultures compared to monocultures, while decreasing the selection for resistant cells.

**Conclusions:** Our results suggest that optimal antiproliferative effects in heterogeneous cancers can be achieved via an integrative therapeutic approach targeting sensitive and resistant cancer cell populations within a tumor, respectively.

## Background

In the past few years, several new therapies have contributed to the treatment of various human cancers. In addition to the classical complex surgical, radio- and chemotherapy, the emergence of novel targeted(1, 2) and immunotherapies(3) resulted in longer progression-free and overall survival(3, 4). In hormone-receptor-positive (HR+), human epidermal growth factor receptor 2-negative (HER2-) breast cancer CDK4/6 inhibitors and mammalian target of rapamycin (mTOR) inhibitors are the most widely used targeted therapies, adding significant benefit to baseline endocrine therapy(4, 5).

A subset of patients receiving targeted therapies observe disease progression(6, 7). Recent progress indicates that tumor heterogeneity and subclonal evolution can be key features contributing to drug resistance(8-11). Following clonal expansion, acquired mutations in cancer cells give rise to different subclones, populations of distinct geno- and phenotypic characteristics and provide a basis for adaptive evolution of the tumor mass(8, 10). In the case of selective pressure, resistant subclones can exhibit a relative proliferative advantage compared to sensitive cells, resulting in resistant cells becoming the predominant subclones, eventually overtaking the entirety of the tumor mass(8). These resistant subclones can be therapy-induced (i.e. they have not been present as a population before the start of therapy); however, a growing body of evidence confirms that in several cases pre-existing resistant subclones are being selected for during the course of treatment(8, 10, 12-14).

Most current standard-of-care therapy regimens are altered only when chemoresistance renders the tumor mass unresponsive to the drug, resulting in progression or relapse(15-17). Previously effective treatments lose their ability to control the tumor burden and because cross-resistance renders several secondary drug classes ineffective, efficacious second-line treatments can be difficult to find(17, 18). Some of these resistance traits include rewiring key pro-proliferative pathways which can create acquired and targetable sensitivities(19).

Therapeutic approaches could benefit from taking into account evolutionary processes in cancer to develop new tools to postpone or overcome drug resistance. Adaptive therapy aims to exploit the changing proliferative advantage between resistant and sensitive cells. This approach succeeds when resistant cells are more fit compared to sensitive cells when drug pressure is on, while when no treatment is present sensitive cells are more fit(20-22). Another approach in treating both sensitive and resistant cells without providing relative proliferative benefit to either cell type is the application of collateral sensitivity. Collateral sensitivity is the acquired vulnerability of a resistant cell against a second drug, which was not applied previously when resistance for the preceding drugs was generated(23, 24). Exploiting collateral sensitivity aims to control the tumor burden through a combination of drugs by targeting sensitive cells with the standard-of-care primary drug while targeting the acquired sensitivities of resistant cells with a secondary drug(17, 23, 24). Recent clinical trials targeting frequent resistance mechanisms up-front revealed a clear advantage over only blocking the primary target in EGFR-mutant non-small-cell lung cancer and BRAF-mutant melanoma(25, 26). In addition to cancer treatments, collateral sensitivities of antibiotic-resistant bacteria are also highly sought after to propose novel, more effective antibacterial treatment regimens in an era of emerging antibiotic resistance(27).

Here, we developed a coculture system to study the effects of collateral sensitivity on the growth of spheroids containing cells sensitive and resistant to CDK4/6 inhibitors. Transcriptional profiling of sensitive and resistant cells revealed druggable acquired vulnerabilities of the resistant cells. By labeling the sensitive and resistant cells with different fluorescent proteins we were able to track their proliferation under drug pressure, mimicking the population dynamics of sensitive and resistant subclones. Our results show that coculture spheroids of sensitive and resistant cells under the selective pressure of ribociclib selects for the ribociclib-resistant cells. Comparing the transcriptional differences of sensitive and resistant cells revealed the downregulation of G2/M checkpoint in resistant cells, upon which collateral sensitivity against the Wee-1 inhibitor adavosertib (AZD1775) was confirmed in the resistant cells. The combination of ribociclib and adavosertib outperformed the antiproliferative effect in the cocultures but not in the monocultures, while also decreasing the selection for resistant cells. Our results promote a phenotype-driven optimization of evolutionary antiproliferative therapy as a model for further assessment in the pre-clinical and clinical setting.

## Methods

### Cell lines and reagents

The estrogen-receptor-positive (ER+), HER2-CAMA-1 breast cancer cell line was maintained in DMEM + 10% FBS + 1% antibiotic-antimycotic solution. CAMA-1 cell line was authenticated by the ATCC cell line authentication service. Cells were continuously treated with ribociclib (Selleck Chemicals, Cat. No: S7440) at 1µM concentration for 1 month followed by 250 nM for 4 months to develop resistance. Ribociclib-resistant CAMA-1 cells (CAMA-1_ribociclib_resistant) were further maintained in complete culture medium supplemented with 250 nM ribociclib. Resistance and collateral sensitivity against adavosertib (Selleck Chemicals, Cat. No.: S1525) were detected by the alteration of the dose-response curve measured using CellTiterGlo Chemoluminescent Kit (Promega Corporation, Cat. No.: G7573). Cell lines were confirmed to be mycoplasma-negative using the Mycoalert PLUS Mycoplasma detection kit (Lonza, Cat. No.: LT07-703).

### Sequencing and bioinformatic analysis

Parental sensitive CAMA-1 and CAMA-1_ribociclib_resistant cell lines were plated 500,000 cells/well in a 6-well plate in triplicates. 24 hours after plating 1 µM ribociclib or vehicle (dimethyl sulfoxide, DMSO) treatment were applied for 12 hours, after which cells were trypsinized, washed and the pellet was frozen at −80C for subsequent RNA isolation. RNA was isolated using the RNeasy Plus Mini Kit (Qiagen, Cat. No.: 74136) following the manufacturer’s protocol.

RNA-seq libraries were prepared using Illumina TruSeq Stranded Total RNA library Prep Ribo-zero Gold following manufacturer’s protocol. Libraries were sequenced with biological triplicates on an Illumina NovaSeq 6000 instrument with 2 × 150 paired-end reads resulting in an average of 25 million reads per sample. Samples were aligned to the human reference genome (hg19) using the STAR (v2.7.0) aligner(28). Transcripts were quantified by RSEM (v1.3.1) followed by differential expression analysis using DESeq2 (v1.26) and GSEA pathways analysis using TPM normalized values and the R packages GSVA (v1.30.0)(29-31). Genes with at least a 2-fold change in expression with FDR < 0.05 were considered statistically significant. Signature scores were generated using the Molecular Signatures Database (v6) Hallmark signature sets. Pathway enrichments with global p-value < 0.05 and FDR < 0.25 were considered statistically significant. Differentially expressed genes were also subjected to pathway analysis regarding the Biocarta pathways using DAVID Bioinformatics Resources(32). In this analysis an FDR-corrected p-value < 0.05 was considered statistically significant.

DNA was isolated from CAMA-1 and CAMA-1_ribociclib_resistant cells using DNeasy Blood & Tissue Kit (Qiagen, Cat. No.: 69504) according to the manufacturer’s protocol. For whole-exome sequencing, libraries were prepared using Agilent SureSelect XT human All Exon v7 following the manufacturer’s protocol. Libraries were sequenced on a NovaSeq 6000 with 2 X150 paired-end reads to a sequencing depth of 285X (CAMA-1) and 300X (CAMA-1_ribociclib_resistant). Reads were trimmed with Trimmomatic prior to alignment to hg19 using BWA-mem(33, 34). Genome Analysis Tool Kit (GATK) best practice guidelines were then followed including the use of Picard and Samtools for PCR duplicate removal and bam manipulation, and GATK for indel realignment and base recalibration(35, 36). Variant calling was performed using an n −1 consensus approach using three somatic variant callers: Mutect2, Strelka and Varscan2(35, 37, 38). This approach was chosen to reduce caller specific false positives. Variant annotations were generated using SNPeff, and Annovar(39, 40). All bioinformatic analysis utilized the BETSY workflow manager(41).

### Lentiviral labeling of sensitive and resistant cells

Lentiviruses incorporating Venus (LeGO-V2) and mCherry (LeGO-C2) fluorescent proteins were generated using Lipofectamine 3000 reagent (Thermo Fisher Scientific) according to the manufacturer’s instructions. LeGO-V2 and LeGO-C2 vectors were gifts from Boris Fehse (Addgene plasmids #27340 and #27339)(42). CAMA-1 and CAMA-1_ribociclib_resistant cell lines were transduced with Venus- and mCherry-containing lentivirus, respectively using reverse transduction, resulting in CAMA-1_V2 and CAMA-1_riboR_C2 cell lines. Briefly, 1 mL of polybrene-containing cell suspension of 200,000 cells were plated in a well of a 6-well plate, where 0.5 ml of viral aliquot was previously dispensed. Cells were incubated for 48 hours at 37°C and 5% CO_2_, after which cells were washed and fresh regular culture medium was applied. Fluorescently labeled cells were selected using fluorescence-activated cell sorting after further subculture of transduced cells to attain homogeneously labeled cell populations.

### Coculture experiments

2000 cells were plated in different proportions (100% CAMA-1_V2, 50% CAMA-1_V2 – 50% CAMA-1_riboR_C2, 100% CAMA-1_riboR_C2) in 96-well round-bottom ultra-low attachment spheroid microplate (Corning, Cat. No.: 4520). 24 hours later, spheroids were washed and fresh medium including treatment drugs was applied. Spheroids were treated for a total of 21 days with imaging and media change been performed at every 4^th^ and 7^th^ day of the week. Imaging was performed using Cytation 5 imager (Biotek Instruments) gathering signal intensity from brightfield, YFP (for Venus fluorescence) and Texas Red (for mCherry fluorescence) channels. Raw data processing and image analysis were performed using Gen5 3.05 software (Biotek Instruments). Briefly, the stitching of 2X2 montage images and Z-projection using focus stacking was performed on raw images followed by spheroid area analysis. On the 21^st^ day of treatment, spheroids were harvested, trypsinized, washed, resuspended in 2 µg/mL DAPI containing flow cytometry buffer (PBS + 5% FBS). Samples were subjected to flow cytometry analysis using Fortessa X20 flow cytometer (BD Biosciences) to assess the relative proportion of sensitive (Venus-labeled) and resistant (mCherry-labeled) cells. Venus fluorescence was excited with 488 nm laser and was separated by a 505 long-pass filter and detected through a 530/30 bandpass filter. mCherry fluorescence was excited with 561 nm laser and was separated by a 595 long-pass filter and detected through a 610/20 bandpass filter. DAPI fluorescence was excited with 355 nm laser and was separated by a 410 nm long-pass filter and detected through a 450/50 bandpass filter. Flow cytometry data were analyzed using FlowJo software (FlowJo, LLC version 10.5.3). Following the exclusion of doublets, fluorescence-positive live cells were analyzed in order to determine the proportion of sensitive and resistant cells within the spheroids. All coculture experiments were performed in triplicates.

### Statistical analysis

Dose-response curves were generated using GraphPad Prism 7.02 software. Differences in dose-response curves were compared using extra sum-of-squares F-test. Differences in spheroid areas and cell proportions were analyzed using Student’s independent samples T-test. Unless otherwise stated a p-value < 0.05 was considered statistically significant.

## Results

### Long-term ribociclib treatment results in ribociclib-resistant cell line

After months of continuous ribociclib treatment, resistance to ribociclib emerged (CAMA-1_ribociclib_resistant cell line), as demonstrated with the different dose-response curves (p<0.0001 Figure 1, Panel A). Short-term ribociclib treatment resulted in 151 and 69 differentially expressed genes in the sensitive (CAMA-1) and resistant (CAMA-1_ribociclib_resistant) cell lines, respectively, with 68 genes downregulated in both cell lines (Figure 1, Panel B-D, Supplementary Tables 1-2). Using both Hallmark and Biocarta pathway gene sets, we find that “CDK regulation of DNA replication” was significantly altered in the resistant cells (untreated vs. ribociclib treated CAMA-1_ribociclib_resistant cells, FDR-corrected p-value: 7.2×10^−6^) while the two significantly dysregulated pathways between untreated and treated sensitive CAMA-1 cells were “CDK regulation of DNA replication” (FDR-corrected p-value: 6.6×10^-10^) and “cyclins and cell cycle regulation” (FDR-corrected p-value: 3.0×,10^-2^), which is consistent with the mode of action of ribociclib.

**Figure 1.**
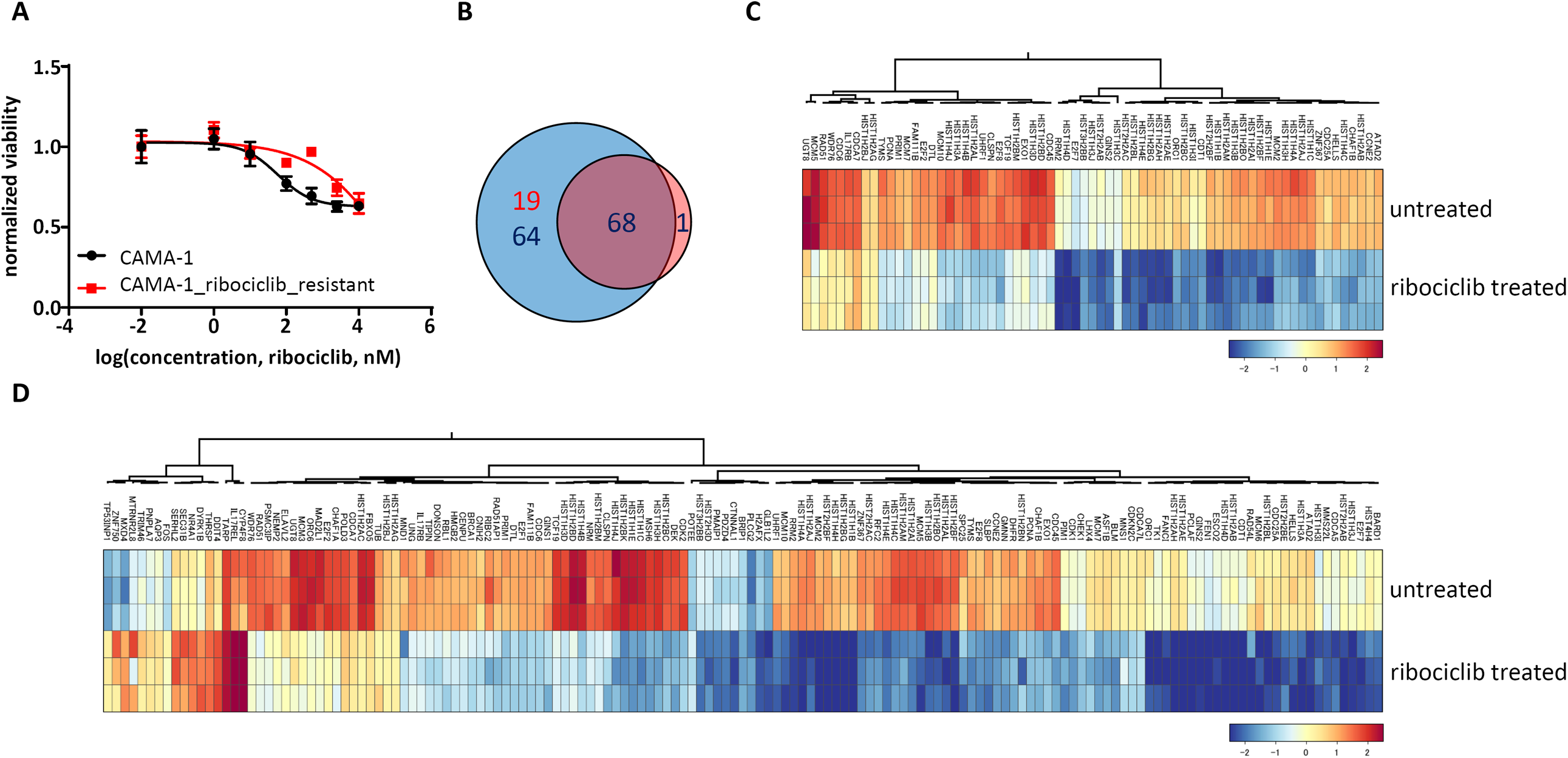
Transcriptional response to ribociclib in sensitive and ribociclib-resistant cells. **Panel A**: Dose-response curves of CAMA-1 and CAMA-1_ribociclib_resistant cells under different concentrations of ribociclib treatment. Cells were treated with increasing concentration of ribociclib for 96 hours, after which viability was measured using CellTiterGlo Chemiluminescent kit. The measured luminescence was normalized to the average of the lowest applied concentration (0.01 nM). Data points show the average of three replicates, error bars show standard deviation if it is larger than the size of the data point. **Panel B**: Venn diagram demonstrating the number of significantly differentially expressed genes in response to 12 hours of 1 µM ribociclib treatment in CAMA-1 and CAMA-1_ribociclib_resistant cells. Blue circle incorporates differentially expressed genes in CAMA-1, while the red circle incorporates differentially expressed genes in CAMA-1_ribociclib_resistant cells. Red numbers demonstrate the number of upregulated, while blue numbers demonstrate the number of downregulated genes in response to ribociclib treatment. **Panel C and D**: Heatmaps demonstrating the expression of significantly differentially expressed genes in response to ribociclib treatment in CAMA-1_ribociclib_resistant (Panel C) and CAMA-1 (Panel D) cells.

We performed whole-exome sequencing on DNA isolated from the sensitive and resistant cells to detect acquired mutations in resistant cells. We found 9 high impact mutations mainly resulting in early stop codon occurrence and 34 moderate mutations resulting in single amino acid changes, while 18 mutations did not change the amino acid sequence of the coded proteins (Table 1). Although no detected mutations are directly associated with resistance against CDK4/6 inhibition, we found several mutations in cancer-associated genes possibly contributing to the acquired resistance against CDK4/6 inhibition.

**Table 1.**
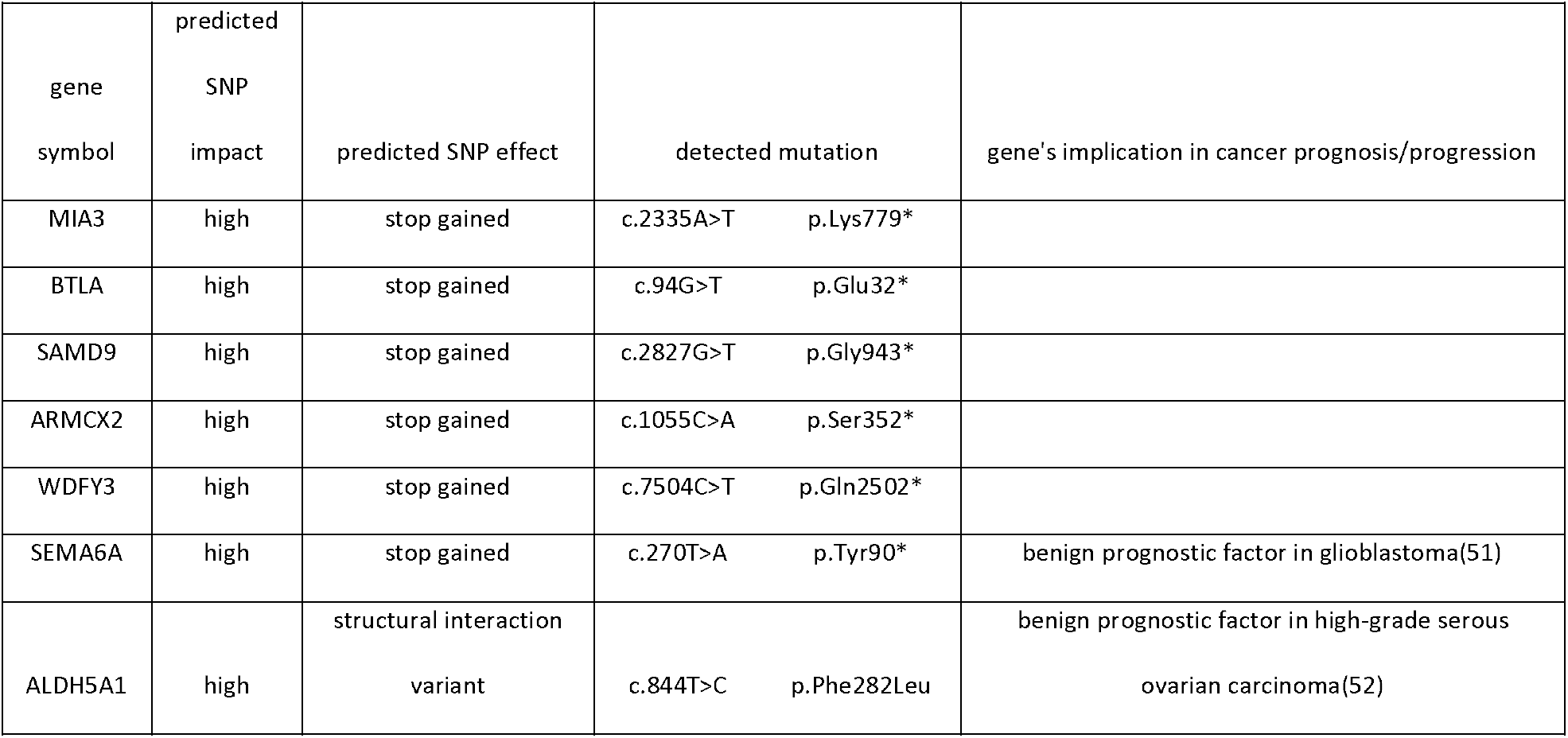
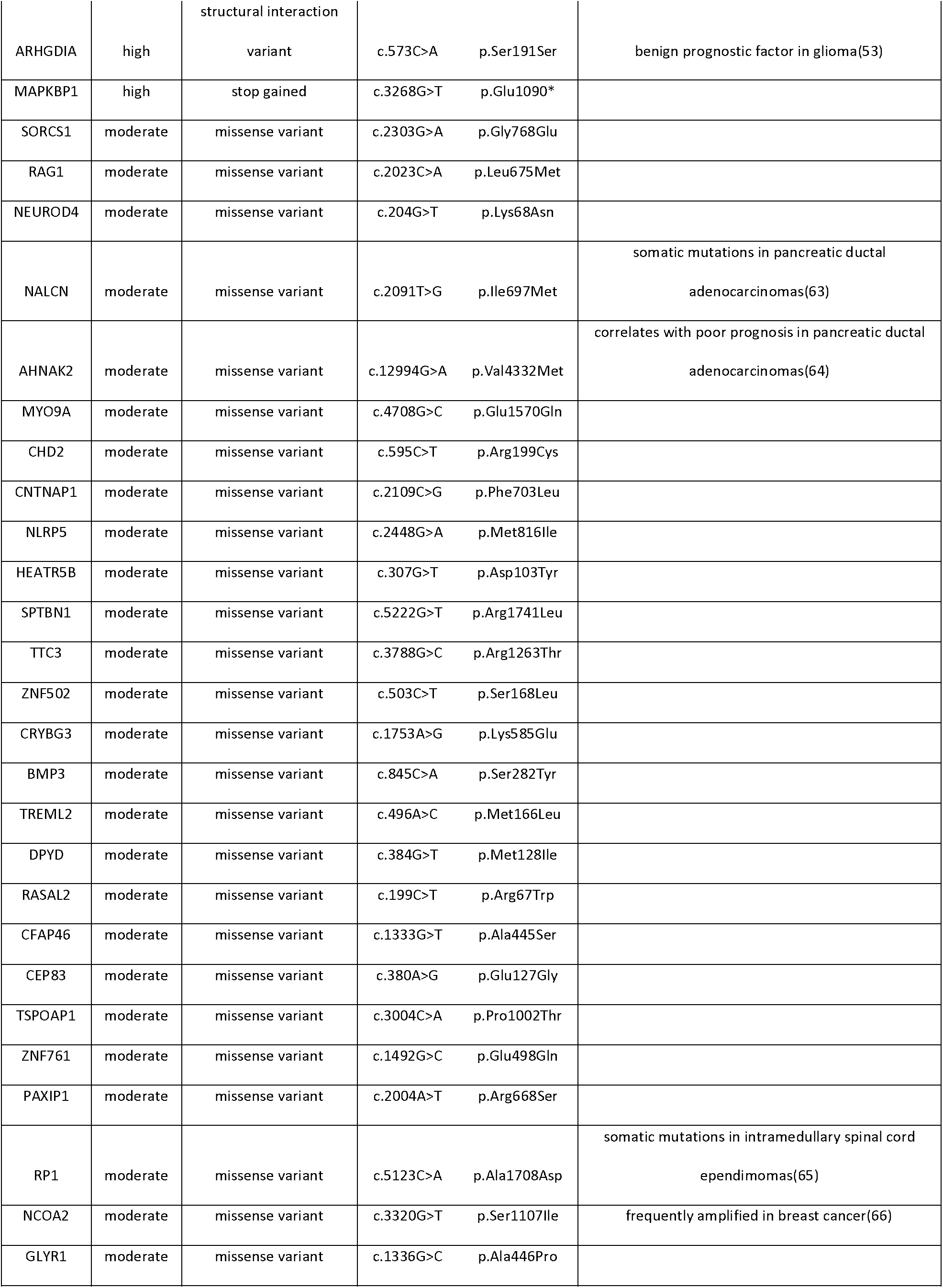
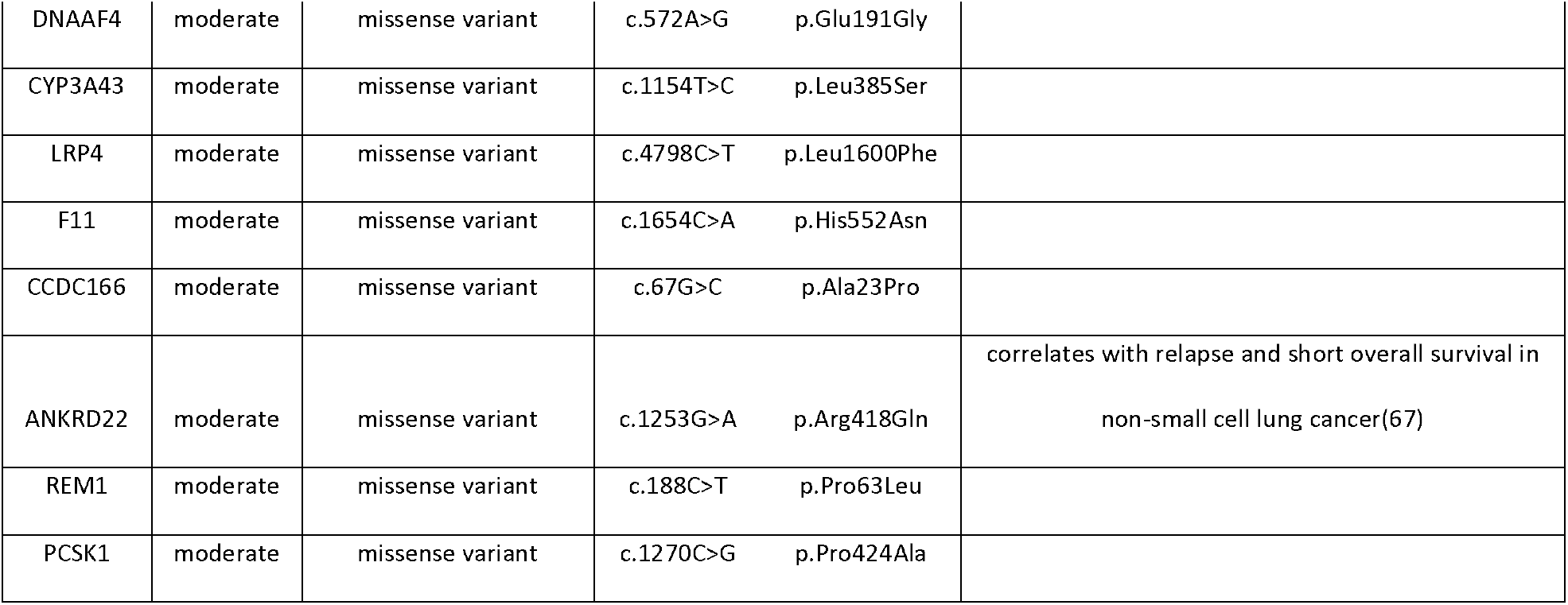
SNPs of high and moderate predicted impact in CAMA-1_ribociclib_resistant cells compared to parental CAMA-1 cells.

| gene symbol | predicted SNP impact | predicted SNP effect | detected mutation | gene's implication in cancer prognosis/progression |
| --- | --- | --- | --- | --- |
| MIA3 | high | stop gained | c.2335A>T p.Lys779* |  |
| BTLA | high | stop gained | c.94G>T p.Glu32* |  |
| SAMD9 | high | stop gained | c.2827G>T p.Gly943* |  |
| ARMCX2 | high | stop gained | c.1055C>A p.Ser352* |  |
| WDFY3 | high | stop gained | c.7504C>T p.Gln2502* |  |
| SEMA6A | high | stop gained | c.270T>A p.Tyr90* | benign prognostic factor in glioblastoma(51) |
| ALDH5A1 | high | structural interaction variant | c.844T>C p.Phe282Leu | benign prognostic factor in high-grade serous ovarian carcinoma(52) |

|  |  |  |  |  |  |
| --- | --- | --- | --- | --- | --- |
| ARHGDIA | high | structural interaction<br>variant | c.573C>A | p.Ser191Ser | benign prognostic factor in glioma(53) |
| MAPKBP1 | high | stop gained | c.3268G>T | p.Glu1090* |  |
| SORCS1 | moderate | missense variant | c.2303G>A | p.Gly768Glu |  |
| RAG1 | moderate | missense variant | c.2023C>A | p.Leu675Met |  |
| NEUROD4 | moderate | missense variant | c.204G>T | p.Lys68Asn |  |
| NALCN | moderate | missense variant | c.2091T>G | p.Ile697Met | somatic mutations in pancreatic ductal<br>adenocarcinomas(63) |
| AHNAK2 | moderate | missense variant | c.12994G>A | p.Val4332Met | correlates with poor prognosis in pancreatic ductal<br>adenocarcinomas(64) |
| MYO9A | moderate | missense variant | c.4708G>C | p.Glu1570Gln |  |
| CHD2 | moderate | missense variant | c.595C>T | p.Arg199Cys |  |
| CNTNAP1 | moderate | missense variant | c.2109C>G | p.Phe703Leu |  |
| NLRP5 | moderate | missense variant | c.2448G>A | p.Met816Ile |  |
| HEATR5B | moderate | missense variant | c.307G>T | p.Asp103Tyr |  |
| SPTBN1 | moderate | missense variant | c.5222G>T | p.Arg1741Leu |  |
| TTC3 | moderate | missense variant | c.3788G>C | p.Arg1263Thr |  |
| ZNF502 | moderate | missense variant | c.503C>T | p.Ser168Leu |  |
| CRYBG3 | moderate | missense variant | c.1753A>G | p.Lys585Glu |  |
| BMP3 | moderate | missense variant | c.845C>A | p.Ser282Tyr |  |
| TREML2 | moderate | missense variant | c.496A>C | p.Met166Leu |  |
| DPYD | moderate | missense variant | c.384G>T | p.Met128Ile |  |
| RASAL2 | moderate | missense variant | c.199C>T | p.Arg67Trp |  |
| CFAP46 | moderate | missense variant | c.1333G>T | p.Ala445Ser |  |
| CEP83 | moderate | missense variant | c.380A>G | p.Glu127Gly |  |
| TSPOAP1 | moderate | missense variant | c.3004C>A | p.Pro1002Thr |  |
| ZNF761 | moderate | missense variant | c.1492G>C | p.Glu498Gln |  |
| PAXIP1 | moderate | missense variant | c.2004A>T | p.Arg668Ser |  |
| RP1 | moderate | missense variant | c.5123C>A | p.Ala1708Asp | somatic mutations in intramedullary spinal cord<br>ependimomas(65) |
| NCOA2 | moderate | missense variant | c.3320G>T | p.Ser1107Ile | frequently amplified in breast cancer(66) |
| GLYR1 | moderate | missense variant | c.1336G>C | p.Ala446Pro |  |
| DNAAF4 | moderate | missense variant | c.572A>G | p.Glu191Gly |  |
| CYP3A43 | moderate | missense variant | c.1154T>C | p.Leu385Ser |  |
| LRP4 | moderate | missense variant | c.4798C>T | p.Leu1600Phe |  |
| F11 | moderate | missense variant | c.1654C>A | p.His552Asn |  |
| CCDC166 | moderate | missense variant | c.67G>C | p.Ala23Pro |  |
| ANKRD22 | moderate | missense variant | c.1253G>A | p.Arg418Gln | correlates with relapse and short overall survival in<br>non-small cell lung cancer(67) |
| REM1 | moderate | missense variant | c.188C>T | p.Pro63Leu |  |
| PCSK1 | moderate | missense variant | c.1270C>G | p.Pro424Ala |  |

### Ribociclib treatment selects for resistant cells in mixed cultures of sensitive and resistant cells

To discriminate sensitive and resistant cells in a coculture system, cell lines were labeled using lentiviral gene transfer with Venus and mCherry fluorescent proteins, respectively. Positively labeled cells were sorted using fluorescence-activated cell sorting to retain the homogenously labeled populations and were further subcultured to be utilized in the spheroid experiments. To analyze the long term effects of ribociclib treatment in 3D cocultures of sensitive and resistant cells, 21-day-long experiments using spheroids of different compositions (100% sensitive, 50% sensitive – 50% resistant, 100% resistant) were initiated (Figure 2, Panel A).

**Figure 2.**
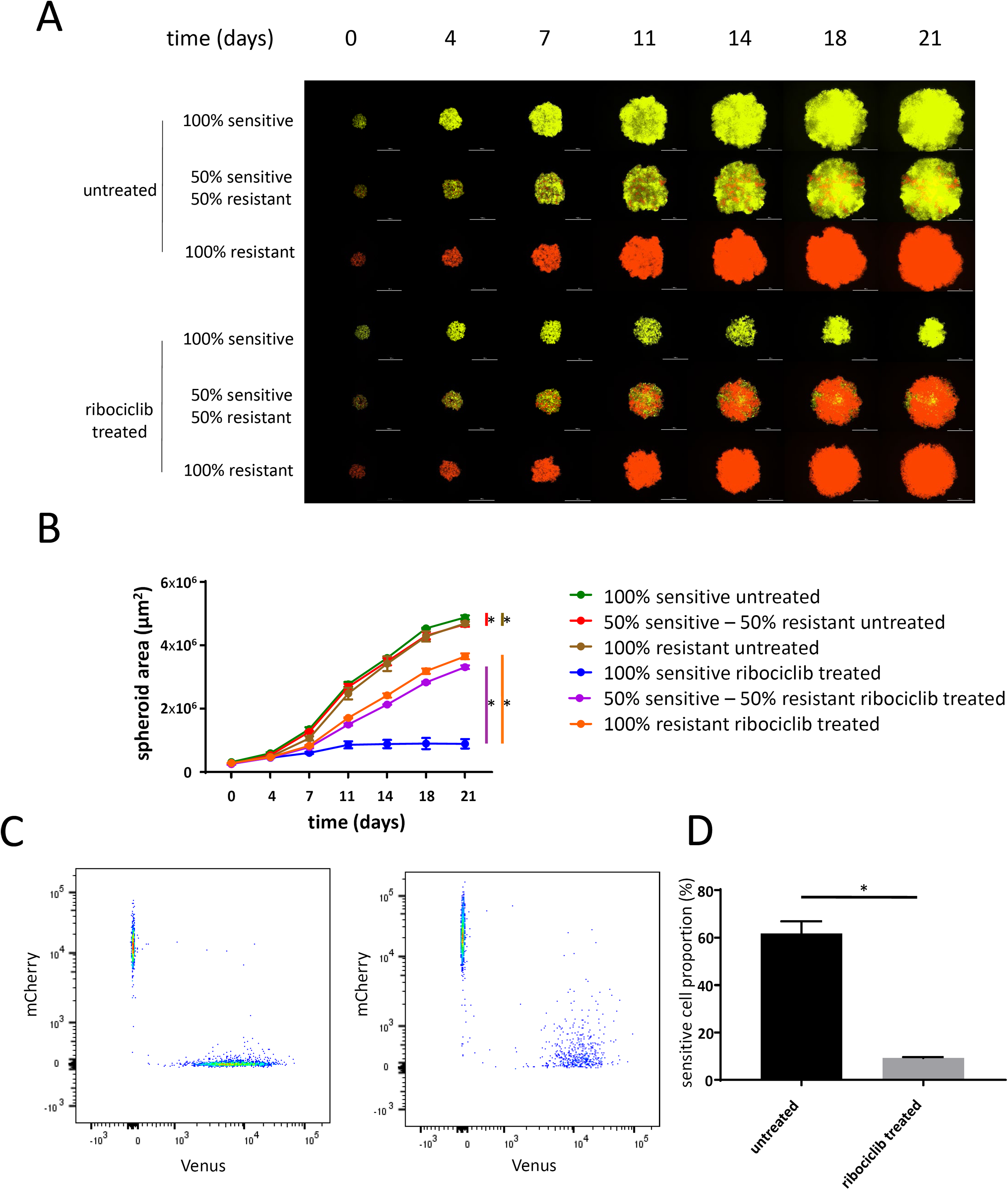
Proliferation of mono- and coculture of sensitive and resistant cells under ribociclib treatment. **Panel A**: Representative images of time course spheroid growth of spheroids of different compositions. Images present the merged images of the Venus and mCherry channels. White bars correspond to 1000 µm. **Panel B**: Time course growth of spheroids presented on Panel A. Data points show average spheroid areas of three replicates, error bars show standard deviation if it is larger than the size of the data point. Vertical lines represent statistical comparisons of the spheroid area between untreated sensitive vs. mixed (red), untreated sensitive and untreated resistant (brown), treated sensitive and mixed (purple) and treated sensitive and resistant (orange) spheroids on Day 21. Asterisks mark statistical significance (p<0.05). Further statistical analysis including all time points and raw data are presented in Supplementary Table 1. **Panel C**: Representative FACS analysis of mixed spheroids on Day 21 on untreated and ribociclib treated spheroids initiated in a composition of 50% sensitive and 50% resistant cells. Samples are identical to the mixed samples on Panel A (second and fifth rows). **Panel D**: Proportion of sensitive cells in untreated and ribociclib treated cocultured spheroids based on FACS analysis. Bars show the average of three replicates, error bars show standard deviation. Asterisk marks statistical significance (p<0.05).

Under no treatment, the growth of all spheroids was remarkable, reaching a 15-17-fold increase in spheroid area during the 21-day treatment (Figure 2, Panel B, Supplementary Table 3). In the coculture setting, sensitive cells exhibited a relative proliferative advantage, as the FACS analysis confirmed that the majority of cells were Venus-labeled sensitive cells (Figure 2, Panels C-D). Continuous ribociclib treatment of 400 nM principally inhibited the growth of the spheroids that were started with 100% sensitive cells (only 3-fold increase in spheroid area over 3 weeks). In contrast, treatment only had a modest effect on the growth of spheroids that were started with 100% resistant cells (12-fold increase in spheroid area over 3 weeks). Resistant cells proliferated more effectively under drug pressure in the coculture as well, resulting in a spheroid area 10% smaller than the treated 100% resistant monoculture. Flow cytometry analysis confirmed that ribociclib treatment selected for the resistant cells (Figure 2, Panels C-D).

### Transcriptional profiling of sensitive and resistant cells reveal key acquired vulnerability of resistant cells

To optimize the treatment of mixed spheroids of sensitive and resistant cells, we aimed to detect acquired vulnerabilities in the phenotype of the resistant cells, upon which collateral sensitivity of potential combination drugs can be applied. We compared the transcriptional profiles of untreated sensitive and resistant cells (untreated CAMA-1 vs. untreated CAMA-1_ribociclib_resistant) and found that 625 genes were significantly dysregulated (fold change > 2 and false discovery rate < 0.05, Supplementary Figure 1, Supplementary Table 4). Among these genes, several cell cycle (CDK6, RUNX2, CCNB3) and antiapoptotic (BCL2) genes were found to be overexpressed in resistant compared to sensitive cells. Pathway analysis using the Hallmark pathways provided 7 pathways significantly dysregulated between sensitive and resistant cells (Figure 3, Panel A-B, Supplementary Figure 2, Supplementary Table 5).

**Figure 3.**
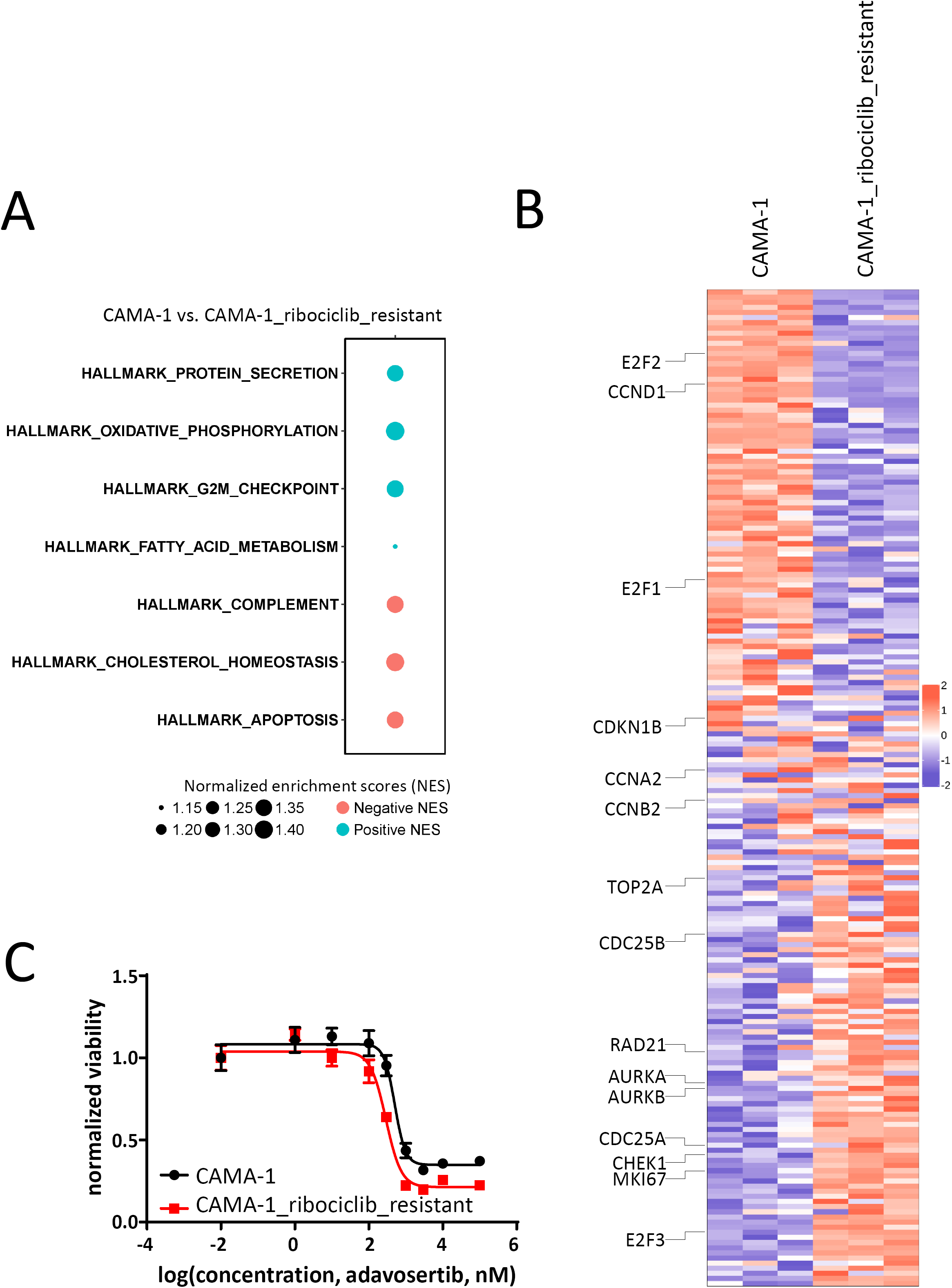
Comparing the transcriptional program of CAMA-1 and CAMA-1_ribociclib_resistant cells reveals collateral sensitivity to Wee-1 inhibition. **Panel A**: Schematic representation of significantly altered Hallmark pathways in between untreated CAMA-1 and CAMA-1_ribociclib_resistant cell lines. Positive normalized enrichment scores (NES) corresponds to enriched pathways in CAMA-1 (blue circles), while pathways with negative NES values (red circles) are enriched in CAMA-1_ribociclib_resistant cells. Additional enrichment scores for Hallmark pathways can be found in Supplementary Table 5. **Panel B**: Heatmap of all genes included in the Hallmark G2/M pathway in untreated CAMA-1 and CAMA-1_ribociclib_resistant cells. Representative genes of the pathway are labeled. Heatmap with all genes labeled can be found in Supplementary Figure 2. **Panel C**: Dose-response curves of CAMA-1 and CAMA-1_ribociclib_resistant cells under different concentrations of Wee-1 inhibitor adavosertib treatment. Cells were treated with increasing concentration of adavosertib for 96 hours, after which viability was measured using CellTiterGlo Chemiluminescent kit. The measured luminescence was normalized to the average of the lowest applied concentration (0.01 nM). Data points show the average of three replicates, error bars show standard deviation if it is larger than the size of the data point.

The loss of the G2/M checkpoint renders cells more susceptible to mitotic catastrophe in the absence of sufficient quality control. Since the CAMA-1_ribociclib_resistant cells are more resistant to the G1 arrest caused by ribociclib and quality control at the G2/M checkpoint is also diminished in these cells based on the transcriptional profiling, Wee-1 inhibition stimulating resistant cells to enter mitotic catastrophe can be more toxic to these cells. Dose-response experiments with the Wee-1-inhibitor, adavosertib, on sensitive and resistant cells showed the IC50 value for adavosertib dropping by 42% (504.2 nM in CAMA-1 cells compared to 291.1 nM in CAMA-1_riboR cells, p-value: 0.0497), confirming the acquired sensitivity of resistant cells against adavosertib (Figure 3, Panel C).

### Leveraging collateral sensitivity controls spheroid growth without selecting for resistant cells

In an effort to overcome the selection of resistant cells in mixed spheroids, while still taking advantage of the antiproliferative effects of ribociclib treatment, we designed a follow-up experiment leveraging the collateral sensitivity of resistant cells in response to adavosertib. The same coculture system was used to investigate the long-term effects of to ribociclib, adavosertib and the combination of these two drugs. While the results of ribociclib treatment replicated our previous experiment, long-term adavosertib treatment proved to be more effective on resistant cells, having only a slight antiproliferative effect on sensitive and mixed spheroids (Figure 4, Panels A and B, Supplementary Table 6). In line with the acquired adavosertib sensitivity in resistant cells, the proportion of sensitive cells was higher in adavosertib treated mixed spheroids, even compared to untreated control mixed spheroids, although no drug pressure also selected for the sensitive cells (Figure 4, Panels C and D). The combination of ribociclib and adavosertib achieved the strongest growth-limiting effect in the spheroids of various composition (100% sensitive, 50% sensitive – 50% resistant, 100% resistant); however, it only added additional amount of growth inhibition compared to single-agent therapy in the case of the mixed spheroids. The growth of 100% sensitive spheroids under ribociclib pressure with or without the addition of adavosertib as well as the growth of 100% resistant spheroids under adavosertib pressure with or without ribociclib was quite indistinguishable, concluding that the combination adds considerable antiproliferative effect only in the mixed spheroids. The combination treatment also resulted in a larger proportion of sensitive cells compared to ribociclib mono-treatment as it selected for neither the sensitive nor the resistant cells.

**Figure 4.**
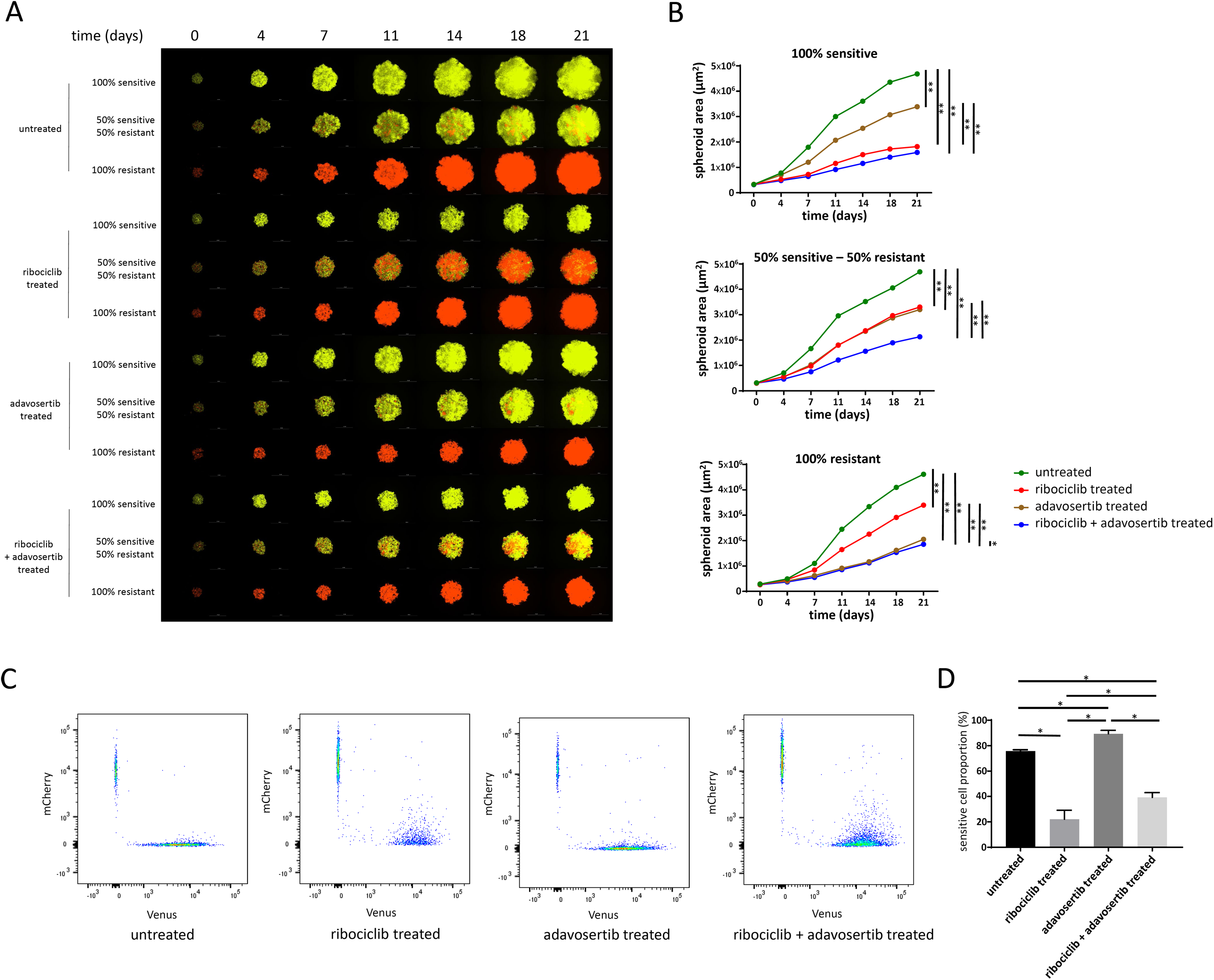
Effects of individual and combination treatments of ribociclib and adavosertib on spheroid growth and composition. **Panel A**: Representative images of time course spheroid growth of spheroids of different compositions. Images present the merged images of the Venus and mCherry channels. White bars correspond to 1000 µm. **Panel B**: Time course growth of spheroids presented on Panel A. Data points show average spheroid areas of three replicates, error bars show standard deviation if it is larger than the size of the data point. Vertical lines represent statistical comparisons of the spheroid area between spheroids on Day 21. Asterisks mark statistical significance (* p<0.05, ** p<0.01). Further statistical analysis including all time points and raw data are presented in Supplementary Table 3. **Panel C**: Representative FACS analysis of mixed spheroids on Day 21 on untreated, ribociclib, adavosertib and ribociclib + adavosertib treated spheroids initiated in a composition of 50% sensitive and 50% resistant cells. Samples are identical to the mixed samples on Panel A (second, fifth, eighth and eleventh rows). **Panel D**: Proportion of sensitive cells under different treatments based on FACS analysis. Bars show the average of three replicates, error bars show standard deviation. Asterisks mark statistical significance (p<0.05).

## Discussion

Leveraging collateral sensitivity aims to control the growth of the tumor mass for a longer period by maintaining a delicate balance between sensitive and resistant cells without providing an unequivocal advantage to either cell type. (17, 23). Since rapid advancements in single-cell sequencing technologies enable us to track each cancer’s subclonal architecture throughout time and treatments(8, 10), personalized clinical management of various tumors in the future might reside on the observed phenotypic differences of resistant subclones and their druggable acquired vulnerabilities.

In the present work, we aimed to create a 3D spheroid system as an in vitro model of a heterogeneous tumor that incorporates both sensitive and resistant cells, followed by analysis of the antiproliferative relevance of specifically targeting both cell types. 3D spheroids recapitulate more closely the transcriptional program of in situ tumor cells compared to monolayer cultures and thus represent an emerging tool to better mimic the in vivo phenotype of tumors affected by cell-cell, cell-matrix interactions(43, 44).

As CDK4/6 inhibitors including ribociclib, abemaciclib, and palbociclib start to occupy a key role in the treatment of HR+, HER2-breast cancer(1), our interest focused on creating a ribociclib-resistant cell line from the widely used and characterized ER+, HER2-CAMA-1 cell line(45, 46). Long-term exposure of ribociclib resulted in CAMA-1_ribociclib_resistant, a ribociclib-resistant CAMA-1 cell line. Sixty-seven and 151 transcripts were differentially expressed in response to short-term ribociclib treatment in the resistant and the sensitive cell lines, respectively. Although many fewer genes were dysregulated in response to ribociclib treatment in the resistant cell line, Biocarta pathway analysis on the dysregulated genes revealed an association with “CDK regulation of DNA replication”. The pathway analysis, as well as the abundance of key members of the cell cycle-dependent transcriptional program (members of the E2F transcription factor family, RRM2, cyclins, MCM2)(47, 48) underlined the clear cell cycle arrest ribociclib achieved in both cell lines. By comparing the baseline transcriptional profile of sensitive and resistant cells, we found CDK6 overexpression in resistant compared to sensitive cells which potentially provides a significant contribution to ribociclib resistance. A previous study also found that CDK6 overexpression is responsible for acquired resistance to CDK4/6 inhibitor abemaciclib in the in vitro setting(49). The overexpression of additional key cell cycle regulators (RUNX2, CCNB3) also may facilitate cell cycle progression in response to CDK4/6 inhibition(50). On the DNA level, we found several acquired mutations in the resistant cells. Although no specific mutated gene can be directly linked to resistance against CDK4/6 inhibitors, high impact mutations in several benign prognostic factors (SEMA6A, ALDH5A1, ARHGDIA) might also contribute to unrestricted proliferation even under drug pressure(51-53).

With the successful creation of the ribociclib resistant cell line and by labeling the cell lines with different fluorescent proteins using lentiviral gene transfer (Venus for CAMA-1 and mCherry for CAMA-1_ribociclib_resistant cell lines), we were able to coculture and track the proliferation of sensitive and resistant cells under different treatments, resulting in a model system resembling heterogeneous tumors. If left to proliferate without drug treatment, sensitive cells enjoyed a proliferative advantage over resistant cells, confirming previous assumptions and observations related to the fitness cost of resistance(16, 54). Long-term ribociclib treatment, however, selected for resistant cells in the coculture setting, as expected. It is also important to note that the growth pattern and the final size of the coculture are only slightly smaller than the 100% resistant monoculture. This reflects the growth of a heterogeneous tumor during which a resistant subclone takes over the tumor mass, diminishing the effect of the primary drug, ribociclib(55).

To overcome the relative proliferative advantage of resistant cells in the coculture setting, and to control the growth of these mixed spheroids more efficiently, we aimed to target sensitive and resistant cells with different drugs. Since sensitive cells responded well to ribociclib, we wanted to combine ribociclib with a secondary drug against which resistant cells developed collateral sensitivity(17). By comparing untreated transcriptional profiles of sensitive and resistant cells, we found that the Hallmark pathway “G2/M checkpoint” was downregulated in resistant cells, which elucidates a potential druggable phenotypic vulnerability. G2/M checkpoint is a major quality control checkpoint during cell cycle progression(56, 57). In the case of inefficient DNA replication, DNA damage response pathways are activated resulting in G2-M arrest. The loss of this quality control checkpoint renders cells with suboptimal cell cycle progression more susceptible to enter mitosis which results in mitotic catastrophe and apoptosis(58). Apoptosis, on the pathway level and the key antiapoptotic regulator BCL2 on the gene level, were both found to be upregulated in resistant cells. This constellation signals apoptosis induction due to continuous ribociclib pressure which is effectively neutralized by elevated BCL2 levels, resulting in the survival of resistant cells under ribociclib pressure(59).

Adavosertib, an inhibitor of the key G2/M checkpoint regulator Wee-1 kinase, was developed to facilitate mitotic catastrophe and subsequent apoptosis in cancer cells where genome integrity primarily relies on a maintained G2/M checkpoint(60-62). We hypothesized that since the G2/M checkpoint is diminished in resistant cells and BCL2 overexpression protects these cells against apoptosis, certain concentrations of adavosertib might achieve a higher antiproliferative effect on resistant compared to sensitive cell lines. Dose-response experiments in sensitive and resistant cell lines confirmed the collateral sensitivity of resistant cells with regard to adavosertib, prompting us for further examination in the long-term coculture system.

Ribociclib mono-treatment confirmed our previous results showing optimal growth inhibitory effect in 100% sensitive spheroids while selecting for the resistant cells in the mixed spheroids. Accordingly, with the acquired sensitivity of the resistant cells, adavosertib mono-treatment had a larger antiproliferative effect on the resistant cells compared to the sensitive cells and selected for the sensitive cells in the mixed setting. Although growth inhibitory effects of the mono-treatments, with the more effective drug on each monoculture (ribociclib for sensitive, adavosertib for resistant), were indistinguishable compared to the combination treatment, combining the two drugs added a considerable antiproliferative effect in the mixed spheroids. This is in line with earlier proposals that controlling the growth of heterogeneous cancers is optimal if all subclones are specifically targeted(8, 17). Additionally, as neither cell type enjoyed a relative proliferative advantage over each other, the spheroids remained highly heterogeneous, keeping a delicate balance between sensitive and resistant cells(20), with the possibility that further treatment with this combination might be durable.

It is important to underline the limitations of our study. Our study is not able to directly translate to in vivo behavior of heterogeneous tumor proliferation, rather it represents a clinically relevant in vitro model system.

## Conclusions

We created an in vitro 3D spheroid coculture system modeling tumor heterogeneity dynamics with respect to sensitive and resistant cells towards the primary CDK4/6-inhibitor treatment. Following transcriptional profiling to detect acquired, collateral sensitivity of resistant cells, we show that an integrative approach selectively targeting sensitive and resistant cells is needed to optimally restrict spheroid growth of cocultures.

## List of abbreviations

CAMA-1_ribociclib_resistant: ribociclib-resistant CAMA-1 cell line
DAPI: 4′,6-diamidino-2-phenylindole
DMSO: dimethyl sulfoxide
ER+: estrogen-receptor-positive
FBS: fetal bovine serum
FDR: false discovery rate
HER2-: human epidermal growth factor receptor 2-negative
HR+: hormone-receptor-positive
mTOR: mammalian target of rapamycin
PBS: phosphate buffered saline

## Declarations

### Availability of data and materials

The datasets supporting the conclusions of this article are available in the Gene Expression Omnibus repository (https://www.ncbi.nlm.nih.gov/geo/; accession number: GSE143944). Additional datasets supporting the conclusions of this article are included within the article and its additional files.

### Competing interests

The authors declare no competing interests.

### Funding

This research and VKG, JC, RE, PAC, LP, AN, PJM, and AHB have been supported by an NIH NCI U54CA209978 grant from the National Cancer Institute. VKG was also supported by the Hungarian Eotvos Scholarship.

The content is solely the responsibility of the authors and does not necessarily represent the official views of the National Institutes of Health.

### Authors’ contributions

VKG performed most of the experiments, was involved in their analysis and drafted the manuscript. JC, LP, AN performed bioinformatics analysis and contributed to manuscript preparation. RE and PAC contributed to performing experiments. PJM contributed to sequencing analysis and contributed to manuscript preparation. AHB conceived the study, contributed to the design and the manuscript preparation. All authors approved the final version of the manuscript.

## Acknowledgements

The authors would like to acknowledge the professional support of the Analytical Cytometry Core of City of Hope National Medical Center and the High-Throughput Genomics Shared Resource at Huntsman Cancer Institute at the University of Utah.

## Supplementary Figures

**Supplementary Figure 1 – Heatmap demonstrating the expression of significantly differentially expressed genes in CAMA-1 and CAMA-1_ribociclib_resistant cells**

**Supplementary Figure 2 – Extended heatmap of Figure 3, Panel B incorporating gene symbols**

**Supplementary Tables**

**Supplementary Table 1 –** Differentially expressed genes in CAMA-1 cells in response to 1 µM ribociclib treatment for 12 hours. Detailed data from Figure 1, Panel D.

**Supplementary Table 2 –** Differentially expressed genes in CAMA-1_ribociclib_resistant cells in response to 1 µM ribociclib treatment for 12 hours. Detailed data from Figure 1, Panel C.

**Supplementary Table 3 – Spheroid area of spheroids plated in different composition under different treatments**

Raw data of three replicates and statistical analysis in each time point of the summarized data in Figure 2, Panel B are shown here.

**Supplementary Table 4 – Differentially expressed genes between CAMA-1 and CAMA- 1_ribociclib_resistant cells under no treatment**

Detailed data from Supplementary Figure 1.

**Supplementary Table 5 – Pathway enrichment scores for the Hallmark pathways in comparison of untreated CAMA-1 and untreated CAMA-1_ribociclib_resistant cells**

Additional data to Figure 3, Panel A.

**Supplementary Table 6 – Spheroid area of spheroids plated in different composition under different treatments**

Raw data of three replicates and statistical analysis in each time point of the summarized data in Figure 4, Panel B are shown here.

## Notes

### Competing Interest Statement

The authors have declared no competing interest.

## References

1. O’Leary B, Finn RS, Turner NC. Treating cancer with selective CDK4/6 inhibitors. Nat Rev Clin Oncol. 2016;13(7):417–30.

2. Hertzman Johansson C, Egyhazi Brage S. BRAF inhibitors in cancer therapy. Pharmacology & therapeutics. 2014;142(2):176–82.

3. Reck M, Rodriguez-Abreu D, Robinson AG, Hui R, Csoszi T, Fulop A, et al. Pembrolizumab versus Chemotherapy for PD-L1-Positive Non-Small-Cell Lung Cancer. The New England journal of medicine. 2016;375(19):1823–33.

4. Im SA, Lu YS, Bardia A, Harbeck N, Colleoni M, Franke F, et al. Overall Survival with Ribociclib plus Endocrine Therapy in Breast Cancer. The New England journal of medicine. 2019;381(4):307–16.

5. Kornblum N, Zhao F, Manola J, Klein P, Ramaswamy B, Brufsky A, et al. Randomized Phase II Trial of Fulvestrant Plus Everolimus or Placebo in Postmenopausal Women With Hormone Receptor-Positive, Human Epidermal Growth Factor Receptor 2-Negative Metastatic Breast Cancer Resistant to Aromatase Inhibitor Therapy: Results of PrE0102. Journal of clinical oncology: official journal of the American Society of Clinical Oncology. 2018;36(16):1556–63.

6. Cross DA, Ashton SE, Ghiorghiu S, Eberlein C, Nebhan CA, Spitzler PJ, et al. AZD9291, an irreversible EGFR TKI, overcomes T790M-mediated resistance to EGFR inhibitors in lung cancer. Cancer Discov. 2014;4(9):1046–61.

7. Johnson DB, Menzies AM, Zimmer L, Eroglu Z, Ye F, Zhao S, et al. Acquired BRAF inhibitor resistance: A multicenter meta-analysis of the spectrum and frequencies, clinical behaviour, and phenotypic associations of resistance mechanisms. European journal of cancer. 2015;51(18):2792–9.

8. Brady SW, McQuerry JA, Qiao Y, Piccolo SR, Shrestha G, Jenkins DF, et al. Combating subclonal evolution of resistant cancer phenotypes. Nature communications. 2017;8(1):1231.

9. Janiszewska M. The microcosmos of intratumor heterogeneity: the space-time of cancer evolution. Oncogene. 2019.

10. Kim C, Gao R, Sei E, Brandt R, Hartman J, Hatschek T, et al. Chemoresistance Evolution in Triple-Negative Breast Cancer Delineated by Single-Cell Sequencing. Cell. 2018;173(4):879–93 e13.

11. Koren S, Bentires-Alj M. Breast Tumor Heterogeneity: Source of Fitness, Hurdle for Therapy. Molecular cell. 2015;60(4):537–46.

12. Choi S, Henderson MJ, Kwan E, Beesley AH, Sutton R, Bahar AY, et al. Relapse in children with acute lymphoblastic leukemia involving selection of a preexisting drug-resistant subclone. Blood. 2007;110(2):632–9.

13. Kim H, Zheng S, Amini SS, Virk SM, Mikkelsen T, Brat DJ, et al. Whole-genome and multisector exome sequencing of primary and post-treatment glioblastoma reveals patterns of tumor evolution. Genome research. 2015;25(3):316–27.

14. Li B, Brady SW, Ma X, Shen S, Zhang Y, Li Y, et al. Therapy-induced mutations drive the genomic landscape of relapsed acute lymphoblastic leukemia. Blood. 2019.

15. Gatenby RA. A change of strategy in the war on cancer. Nature. 2009;459(7246):508–9.

16. Gatenby RA, Brown J, Vincent T. Lessons from applied ecology: cancer control using an evolutionary double bind. Cancer research. 2009;69(19):7499–502.

17. West JB, Dinh MN, Brown JS, Zhang J, Anderson AR, Gatenby RA. Multidrug Cancer Therapy in Metastatic Castrate-Resistant Prostate Cancer: An Evolution-Based Strategy. Clinical cancer research: an official journal of the American Association for Cancer Research. 2019;25(14):4413–21.

18. Ricci F, Guffanti F, Damia G, Broggini M. Combination of paclitaxel, bevacizumab and MEK162 in second line treatment in platinum-relapsing patient derived ovarian cancer xenografts. Mol Cancer. 2017;16(1):97.

19. Ramos P, Bentires-Alj M. Mechanism-based cancer therapy: resistance to therapy, therapy for resistance. Oncogene. 2015;34(28):3617–26.

20. Bacevic K, Noble R, Soffar A, Wael Ammar O, Boszonyik B, Prieto S, et al. Spatial competition constrains resistance to targeted cancer therapy. Nature communications. 2017;8(1):1995.

21. Gatenby RA, Silva AS, Gillies RJ, Frieden BR. Adaptive therapy. Cancer research. 2009;69(11):4894–903.

22. Zhang J, Cunningham JJ, Brown JS, Gatenby RA. Integrating evolutionary dynamics into treatment of metastatic castrate-resistant prostate cancer. Nature communications. 2017;8(1):1816.

23. Dhawan A, Nichol D, Kinose F, Abazeed ME, Marusyk A, Haura EB, et al. Collateral sensitivity networks reveal evolutionary instability and novel treatment strategies in ALK mutated non-small cell lung cancer. Sci Rep. 2017;7(1):1232.

24. Zhao B, Sedlak JC, Srinivas R, Creixell P, Pritchard JR, Tidor B, et al. Exploiting Temporal Collateral Sensitivity in Tumor Clonal Evolution. Cell. 2016;165(1):234–46.

25. Soria JC, Ohe Y, Vansteenkiste J, Reungwetwattana T, Chewaskulyong B, Lee KH, et al. Osimertinib in Untreated EGFR-Mutated Advanced Non-Small-Cell Lung Cancer. The New England journal of medicine. 2018;378(2):113–25.

26. Long GV, Stroyakovskiy D, Gogas H, Levchenko E, de Braud F, Larkin J, et al. Dabrafenib and trametinib versus dabrafenib and placebo for Val600 BRAF-mutant melanoma: a multicentre, double-blind, phase 3 randomised controlled trial. Lancet. 2015;386(9992):444–51.

27. Lazar V, Martins A, Spohn R, Daruka L, Grezal G, Fekete G, et al. Antibiotic-resistant bacteria show widespread collateral sensitivity to antimicrobial peptides. Nat Microbiol. 2018;3(6):718–31.

28. Dobin A, Davis CA, Schlesinger F, Drenkow J, Zaleski C, Jha S, et al. STAR: ultrafast universal RNA-seq aligner. Bioinformatics. 2013;29(1):15–21.

29. Li B, Dewey CN. RSEM: accurate transcript quantification from RNA-Seq data with or without a reference genome. BMC Bioinformatics. 2011;12:323.

30. Love MI, Huber W, Anders S. Moderated estimation of fold change and dispersion for RNA-seq data with DESeq2. Genome biology. 2014;15(12):550.

31. Hanzelmann S, Castelo R, Guinney J. GSVA: gene set variation analysis for microarray and RNA-seq data. BMC Bioinformatics. 2013;14:7.

32. Huang da W, Sherman BT, Lempicki RA. Systematic and integrative analysis of large gene lists using DAVID bioinformatics resources. Nat Protoc. 2009;4(1):44–57.

33. Bolger AM, Lohse M, Usadel B. Trimmomatic: a flexible trimmer for Illumina sequence data. Bioinformatics. 2014;30(15):2114–20.

34. Li H, Durbin R. Fast and accurate short read alignment with Burrows-Wheeler transform. Bioinformatics. 2009;25(14):1754–60.

35. DePristo MA, Banks E, Poplin R, Garimella KV, Maguire JR, Hartl C, et al. A framework for variation discovery and genotyping using next-generation DNA sequencing data. Nature genetics. 2011;43(5):491–8.

36. Li H, Handsaker B, Wysoker A, Fennell T, Ruan J, Homer N, et al. The Sequence Alignment/Map format and SAMtools. Bioinformatics. 2009;25(16):2078–9.

37. Kim S, Scheffler K, Halpern AL, Bekritsky MA, Noh E, Kallberg M, et al. Strelka2: fast and accurate calling of germline and somatic variants. Nat Methods. 2018;15(8):591–4.

38. Koboldt DC, Zhang Q, Larson DE, Shen D, McLellan MD, Lin L, et al. VarScan 2: somatic mutation and copy number alteration discovery in cancer by exome sequencing. Genome research. 2012;22(3):568–76.

39. Cingolani P, Platts A, Wang le L, Coon M, Nguyen T, Wang L, et al. A program for annotating and predicting the effects of single nucleotide polymorphisms, SnpEff: SNPs in the genome of Drosophila melanogaster strain w1118; iso-2; iso-3. Fly (Austin). 2012;6(2):80–92.

40. Wang K, Li M, Hakonarson H. ANNOVAR: functional annotation of genetic variants from high-throughput sequencing data. Nucleic acids research. 2010;38(16):e164.

41. Chen X, Chang JT. Planning bioinformatics workflows using an expert system. Bioinformatics. 2017;33(8):1210–5.

42. Weber K, Bartsch U, Stocking C, Fehse B. A multicolor panel of novel lentiviral “gene ontology” (LeGO) vectors for functional gene analysis. Molecular therapy: the journal of the American Society of Gene Therapy. 2008;16(4):698–706.

43. LaBarbera DV, Reid BG, Yoo BH. The multicellular tumor spheroid model for high-throughput cancer drug discovery. Expert Opin Drug Discov. 2012;7(9):819–30.

44. Sachs N, de Ligt J, Kopper O, Gogola E, Bounova G, Weeber F, et al. A Living Biobank of Breast Cancer Organoids Captures Disease Heterogeneity. Cell. 2018;172(1-2):373–86 e10.

45. Leung BS, Qureshi S, Leung JS. Response to estrogen by the human mammary carcinoma cell line CAMA-1. Cancer research. 1982;42(12):5060–6.

46. Kytola S, Rummukainen J, Nordgren A, Karhu R, Farnebo F, Isola J, et al. Chromosomal alterations in 15 breast cancer cell lines by comparative genomic hybridization and spectral karyotyping. Genes, chromosomes & cancer. 2000;28(3):308–17.

47. Grolmusz VK, Toth EA, Baghy K, Liko I, Darvasi O, Kovalszky I, et al. Fluorescence activated cell sorting followed by small RNA sequencing reveals stable microRNA expression during cell cycle progression. BMC genomics. 2016;17(1):412.

48. Grolmusz VK, Karászi K, Micsik T, Tóth EA, Mészáros K, Karvaly G, et al. Cell cycle dependent RRM2 may serve as proliferation marker and pharmaceutical target in adrenocortical cancer. American journal of cancer research. 2016;6(9):2041–53.

49. Yang C, Li Z, Bhatt T, Dickler M, Giri D, Scaltriti M, et al. Acquired CDK6 amplification promotes breast cancer resistance to CDK4/6 inhibitors and loss of ER signaling and dependence. Oncogene. 2017;36(16):2255–64.

50. Qiao M, Shapiro P, Fosbrink M, Rus H, Kumar R, Passaniti A. Cell cycle-dependent phosphorylation of the RUNX2 transcription factor by cdc2 regulates endothelial cell proliferation. The Journal of biological chemistry. 2006;281(11):7118–28.

51. Zhao J, Tang H, Zhao H, Che W, Zhang L, Liang P. SEMA6A is a prognostic biomarker in glioblastoma. Tumour biology: the journal of the International Society for Oncodevelopmental Biology and Medicine. 2015;36(11):8333–40.

52. Hilvo M, de Santiago I, Gopalacharyulu P, Schmitt WD, Budczies J, Kuhberg M, et al. Accumulated Metabolites of Hydroxybutyric Acid Serve as Diagnostic and Prognostic Biomarkers of Ovarian High-Grade Serous Carcinomas. Cancer research. 2016;76(4):796–804.

53. Lu W, Wang X, Liu J, He Y, Liang Z, Xia Z, et al. Downregulation of ARHGDIA contributes to human glioma progression through activation of Rho GTPase signaling pathway. Tumour biology: the journal of the International Society for Oncodevelopmental Biology and Medicine. 2016.

54. Gallaher JA, Enriquez-Navas PM, Luddy KA, Gatenby RA, Anderson ARA. Spatial Heterogeneity and Evolutionary Dynamics Modulate Time to Recurrence in Continuous and Adaptive Cancer Therapies. Cancer research. 2018;78(8):2127–39.

55. Herrera-Abreu MT, Palafox M, Asghar U, Rivas MA, Cutts RJ, Garcia-Murillas I, et al. Early Adaptation and Acquired Resistance to CDK4/6 Inhibition in Estrogen Receptor-Positive Breast Cancer. Cancer research. 2016;76(8):2301–13.

56. O’Connell MJ, Walworth NC, Carr AM. The G2-phase DNA-damage checkpoint. Trends in cell biology. 2000;10(7):296–303.

57. Lee JH, Paull TT. Activation and regulation of ATM kinase activity in response to DNA double-strand breaks. Oncogene. 2007;26(56):7741–8.

58. Reinhardt HC, Aslanian AS, Lees JA, Yaffe MB. p53-deficient cells rely on ATM- and ATR-mediated checkpoint signaling through the p38MAPK/MK2 pathway for survival after DNA damage. Cancer cell. 2007;11(2):175–89.

59. Knight T, Luedtke D, Edwards H, Taub JW, Ge Y. A delicate balance - The BCL-2 family and its role in apoptosis, oncogenesis, and cancer therapeutics. Biochemical pharmacology. 2019;162:250–61.

60. Fu S, Wang Y, Keyomarsi K, Meric-Bernstam F, Meric-Bernstein F. Strategic development of AZD1775, a Wee1 kinase inhibitor, for cancer therapy. Expert Opin Investig Drugs. 2018;27(9):741–51.

61. Matheson CJ, Backos DS, Reigan P. Targeting WEE1 Kinase in Cancer. Trends Pharmacol Sci. 2016;37(10):872–81.

62. Xu D, Liang SQ, Yang H, Bruggmann R, Berezowska S, Yang Z, et al. CRISPR screening identifies WEE1 as a combination target for standard chemotherapy in malignant pleural mesothelioma. Mol Cancer Ther. 2019.

63. Biankin AV, Waddell N, Kassahn KS, Gingras MC, Muthuswamy LB, Johns AL, et al. Pancreatic cancer genomes reveal aberrations in axon guidance pathway genes. Nature. 2012;491(7424):399–405.

64. Lu D, Wang J, Shi X, Yue B, Hao J. AHNAK2 is a potential prognostic biomarker in patients with PDAC. Oncotarget. 2017;8(19):31775–84.

65. Zhang M, Iyer RR, Azad TD, Wang Q, Garzon-Muvdi T, Wang J, et al. Genomic Landscape of Intramedullary Spinal Cord Gliomas. Sci Rep. 2019;9(1):18722.

66. Cai M, Liang X, Sun X, Chen H, Dong Y, Wu L, et al. Nuclear Receptor Coactivator 2 Promotes Human Breast Cancer Cell Growth by Positively Regulating the MAPK/ERK Pathway. Front Oncol. 2019;9:164.

67. Yin J, Fu W, Dai L, Jiang Z, Liao H, Chen W, et al. ANKRD22 promotes progression of non-small cell lung cancer through transcriptional up-regulation of E2F1. Sci Rep. 2017;7(1):4430.

