## Supplementary figures and images for "Exploiting collateral sensitivity controls growth of mixed culture of sensitive and resistant cells and decreases selection for resistant cells"

### Supplementary figure 2

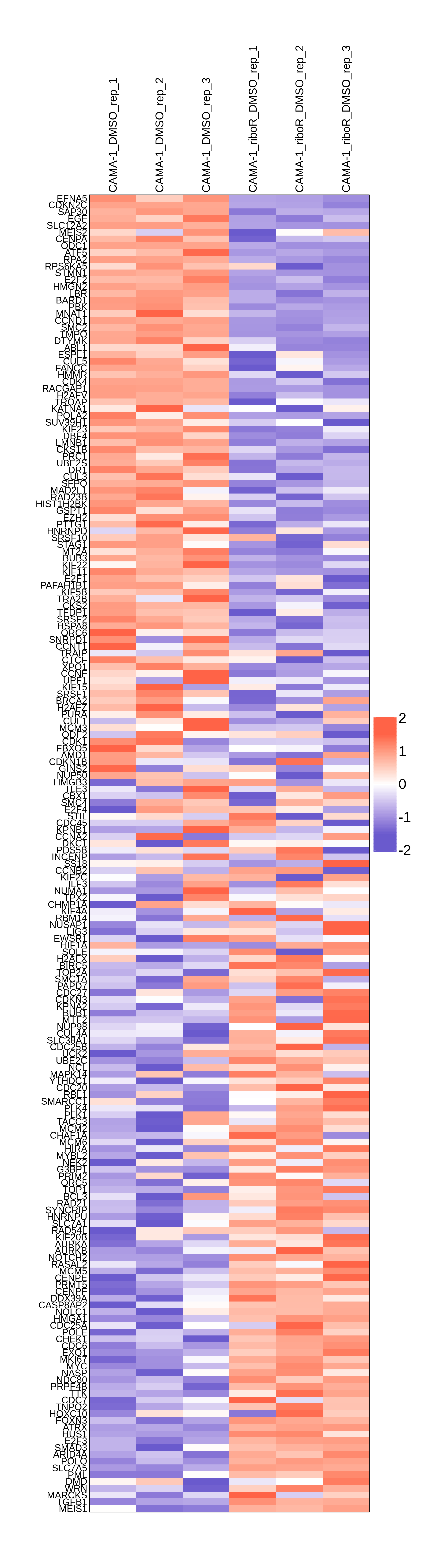
